# Extended spectrum β-lactamase and carbapenemase genes are substantially and sequentially reduced during conveyance and treatment of urban sewage

**DOI:** 10.1101/2020.11.12.379032

**Authors:** Liguan Li, Joseph Nesme, Marcos Quintela-Baluja, Sabela Balboa, Syed Hashsham, Maggie R. Williams, Zhuofeng Yu, Søren J. Sørensen, David W. Graham, Jesús L. Romalde, Arnaud Dechesne, Barth F. Smets

## Abstract

Integrated and quantitative observations of antibiotic resistance genes (ARGs) in urban water systems (UWSs) are lacking. We sampled three UWSs for clinically important extended spectrum β-lactamase (ESBL) and carbapenemase (CP) genes, mobile genetic elements and microbial communities. Sewage – especially from hospitals – carried substantial loads of ESBL and CP genes (10^6^ – 10^7^ per person equivalent), but those loads progressively declined along the UWS, resulting in minimal emissions (10^1^ – 10^4^ copies per person equivalent). Removal was primarily during sewage conveyance (65% ± 36%) rather than within sewage treatment (34% ± 23%). The ARGs clustered in groups based on their persistence; less persistent groups were associated to putative host taxa (especially *Enterobacteriaceae* and *Moraxellaceae*), while more persistent groups appear horizontally transferred as they correlated with mobile genetic elements. This first documentation of a substantial ARG reduction during sewage conveyance provides opportunities for antibiotic resistance management and a caution for sewage-based ARG surveillance.

## Introduction

Antibiotic resistance acquisition by pathogens is one of the biggest global public health challenges, as it increasingly impairs our ability to treat infectious diseases. While the medical and veterinary use of antibiotics is the main cause of the rise of antibiotic resistance, the environment has recently been shown as the single largest reservoir of antibiotic resistance genes (ARGs)^1,2^. The urban water system (UWS) has particularly been suggested as a pathway for antibiotic resistance dissemination through residual waters, as bacteria originating from the human gut are mixed with environmental bacteria, exposed to dynamic conditions and diverse microbial interactions in sewage treatment plants (STP)^3^.

Understanding the fate of antibiotic resistance in the UWS is therefore essential for evaluating its role in ARG dissemination. Raw sewage (i.e., residual water) constitutes the main source of ARGs in the UWS and reflects the load of antibiotic resistance of the served populations. Local, regional and international surveillance studies have shown that influents of urban STP harbor significant abundance and diversity of ARGs with socioeconomic, geographic and seasonal variation^4–6^. Most recently, an integrated surveillance of STP influent in seven European countries revealed ARG profiles that mirrored the north-to-south gradients in clinical antibiotic resistance prevalence across Europe^4^. STP influents often contains sewage not only from domestic origin but also from hospital sources, with contrasting contributions to the total ARG load. Indeed, compared to domestic sewage, hospital sewage has a higher relative load of multi-resistant bacteria and genes, especially clinically important β-lactamase genes^7–11^. It is however not clear to what extent sewage collection and treatment can act as barrier for those clinically important ARGs. Incomplete removal of clinical ARGs by sewage treatment processes may cause further dissemination of antibiotic resistance in the receiving environments^8,12^. Moreover, there is a debate on whether or not separately treating hospital sewage to a higher degree than the rest of sewage would substantially reduce ARG discharge^11^. In addition, the biological treatment units of STPs have been recognized as a location with substantial amount of ARGs and mobile genetic elements (MGEs), with potential for ARG transfer between environmental bacteria and human pathogens^13^.

Realizing that UWSs consist of compartments with very different physical-chemical conditions, efforts to describe ARG dynamics have mainly considered one or a few of these compartments at a time, such as the influent, the biological treatment reactor, and the effluent^4,10,12,14^. However, these efforts fail to capture a full picture of UWS and rigorous studies on the occurrence of ARGs across the different compartments are absent, hampering an integrative understanding of the fate of ARGs from raw sewage, through conveyance, treatment processes, and until final discharge. As sewage travels through the UWS, the microbial communities and their ARGs are diluted, mixed with resident communities^15,16^, exposed to varying environmental conditions and, being sequentially subject to both stochastic (e.g., dispersal and drift) and deterministic selective processes (e.g., by differing nutrient and redox conditions)^17,18^. These processes can lead to the loss of some ARGs if their hosts are driven to extinction, whereas transfer of mobile ARGs can increase their likelihood of persistence, even though the original intestinal host bacteria may die off. It is therefore essential to track ARGs together with community structure and mobility potential to help identify the processes that mitigate or favor antibiotic resistance across the entirety of the UWS. This has not been achieved because most previous studies were based on analysis of ARGs alone and on observations from specific compartments^4,8,12^.

In this study, we systematically investigated the spatiotemporal variation of ARG diversity and abundance across selected, well-characterized UWSs. As single sampling regions/events fail to capture the variability of antibiotic resistance load^4^, we conducted detailed sampling campaigns in parallel in three cities in Denmark (DK), Spain (SP) and the United Kingdom (UK), respectively, with differing ARG occurrence and prescription patterns. However, all cities had modern sewage collection and STP infrastructure. We analyzed both the bacterial communities and 70 clinically important β-lactamase genes, from raw sewage of hospital and residential sources, through different treatment processes in STP, until the final discharge and the receiving river. In all the three countries, we consistently observed significantly greater antibiotic resistance contributions from hospital sources within the UWSs, and a dramatic reduction in ARG abundance during conveyance and treatment. We identified ARG groups with different persistence fate across UWS, which were correlated with the fate of distinct taxonomic groups.

## Results

### Community and ARG profile across countries, compartments and campaigns

We sampled key compartments within three UWSs: Odense (DK), Santiago de Compostela (SP), and Durham (UK) (Fig. 1, Table S1). These compartments were (the two letter code in parenthesis used in all figures): sewage from the city’s hospital (HS); sewage from a residential area (RS); influent sewage to the STPs (IS), after screening and grit removal; primary clarifier (PC) effluent; the main biological treatment reactor (BT), i.e., activated sludge in DK and SP and a biofilter in UK; and secondary clarifier (SC) effluent discharged to the environment in the UK and SP; in DK, a tertiary treatment (TT, multimedia filtration and post aeration) is applied before discharge. Finally, the river receiving the STP effluent was sampled both upstream and downstream of the discharge point (RU and RD, respectively). Additional information on sampling is provided in Supplementary Information.

**Fig. 1.**
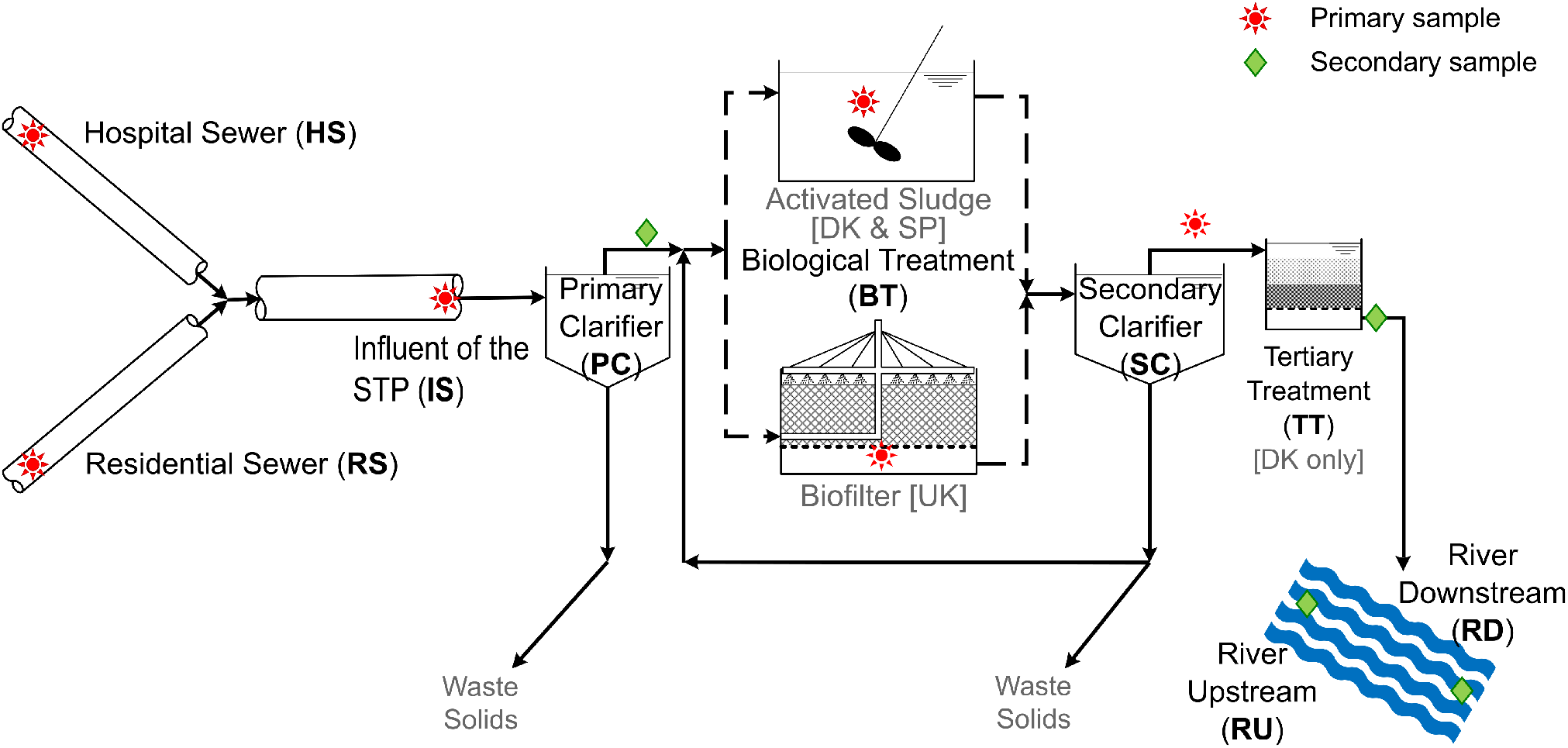
Sampling sites of UWSs in three European cities: Odense (DK), Santiago de Compostela (SP), and Durham (UK). The two letter codes of sampling compartments in parenthesis are used in all figures.

Overall microbial community compositions and ARG profiles were quantified across sample sets (Fig. 2 & 3). The concentration of bacteria across all samples ranged from 4.5 × 10^9^ cells per ml in STP effluent to 3.8 × 10^13^ cells per ml in hospital sewage. Similar compartments dominated shaping community profiles (PERMANOVA, P-value < 0.01), even though samples were collected from three geographically distinct European countries (i.e., DK, SP, UK) over three campaigns (spanning about 1.5 year) (Fig. S1). Clearly, hospital and residential sewage had the most similar microbial community composition (average Bray-Curtis dissimilarity = 0.35), which was also similar to treatment plant influent and the primary clarifier. The dominant taxa in raw sewage were of human-gut origin, with members of *Clostridriales*, *Bacteroidales* and *Enterobacteriales* orders accounting for 37% of the community on average, and even reaching > 50% in 7 out of 18 sewage samples from the three countries. These enteric bacteria were lost in the influent and primary clarifier, where the communities became mainly composed of *Campylobacterales*, *β-proteobacteriales*, and *Pseudomonadales*, together accounting 48% on average. The community structure shift across compartments followed a similar pattern in the three countries until the biological treatment compartment. There, the microbial community profile of DK and SP (mainly composed by *β-proteobacteriales*, *Chitinophagales*, *Micrococcales*) significantly differed from that of the UK (mainly composed by *Pseudomonadales*, *β-proteobacteriales*, *Flavobacteriales*) where a biofilter-based process was applied, while the STP in DK and SP employed a suspended-growth process. The concentration of several taxa listed above increased in biological treatment compartment, before being substantially removed by secondary clarification.

**Fig. 2.**
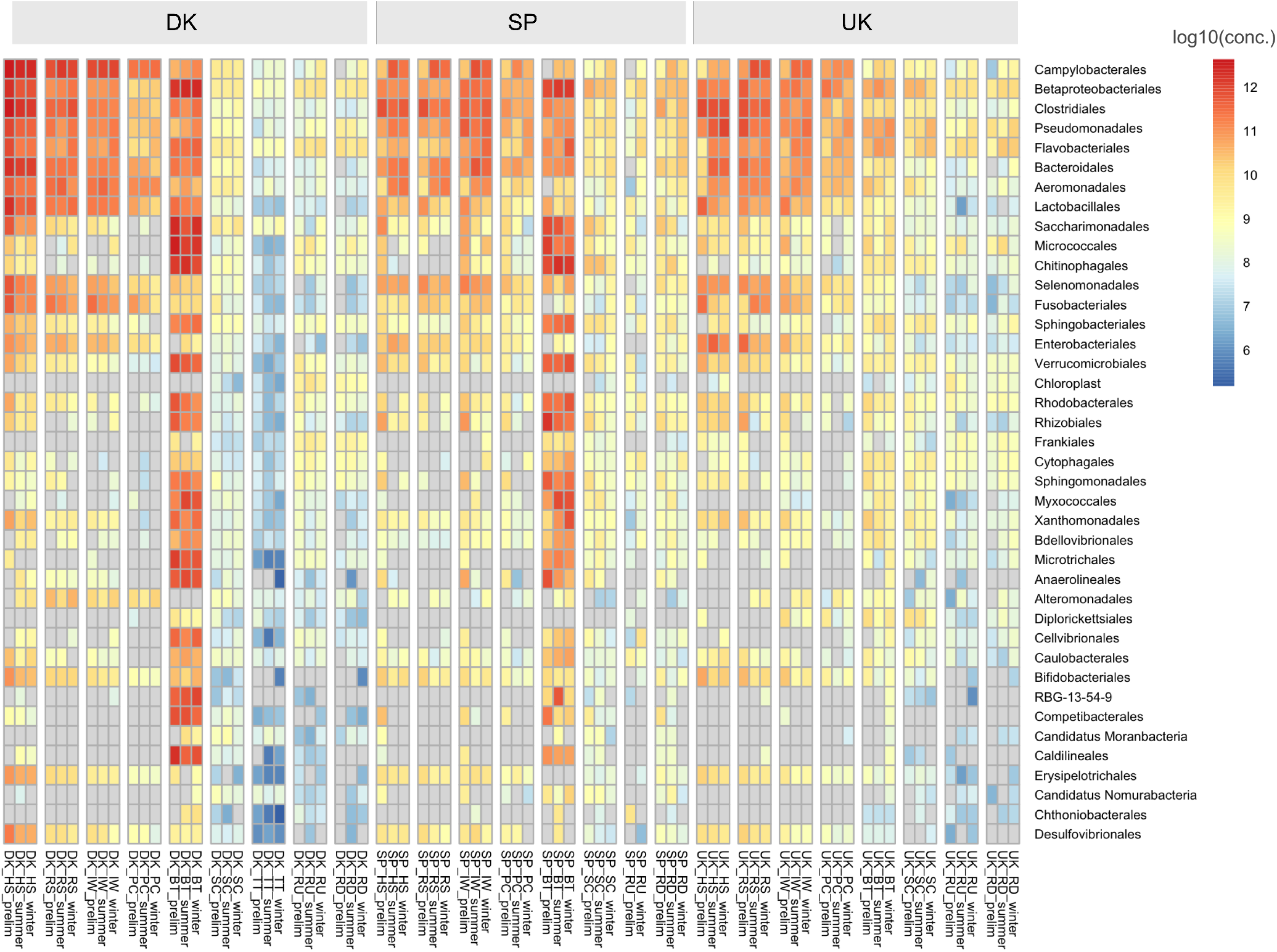
Concentration heatmap of the 40 most abundant taxa at order level across the three countries. The rows are 40 orders arranged from high to low average abundance across all samples. Each column indicates ARG concentration in specific country and UWS compartment averaged by sampling days in the campaign. For each country, columns are ordered by compartment sequence (HS-RS-IW-PC-BT-SC-TT-RU-RD) through UWS of the country. Concentration (cells per ml) is log10 transformed. Color scale from blue to red indicates from low to high concentration, and gray indicates not detected.

**Fig. 3.**
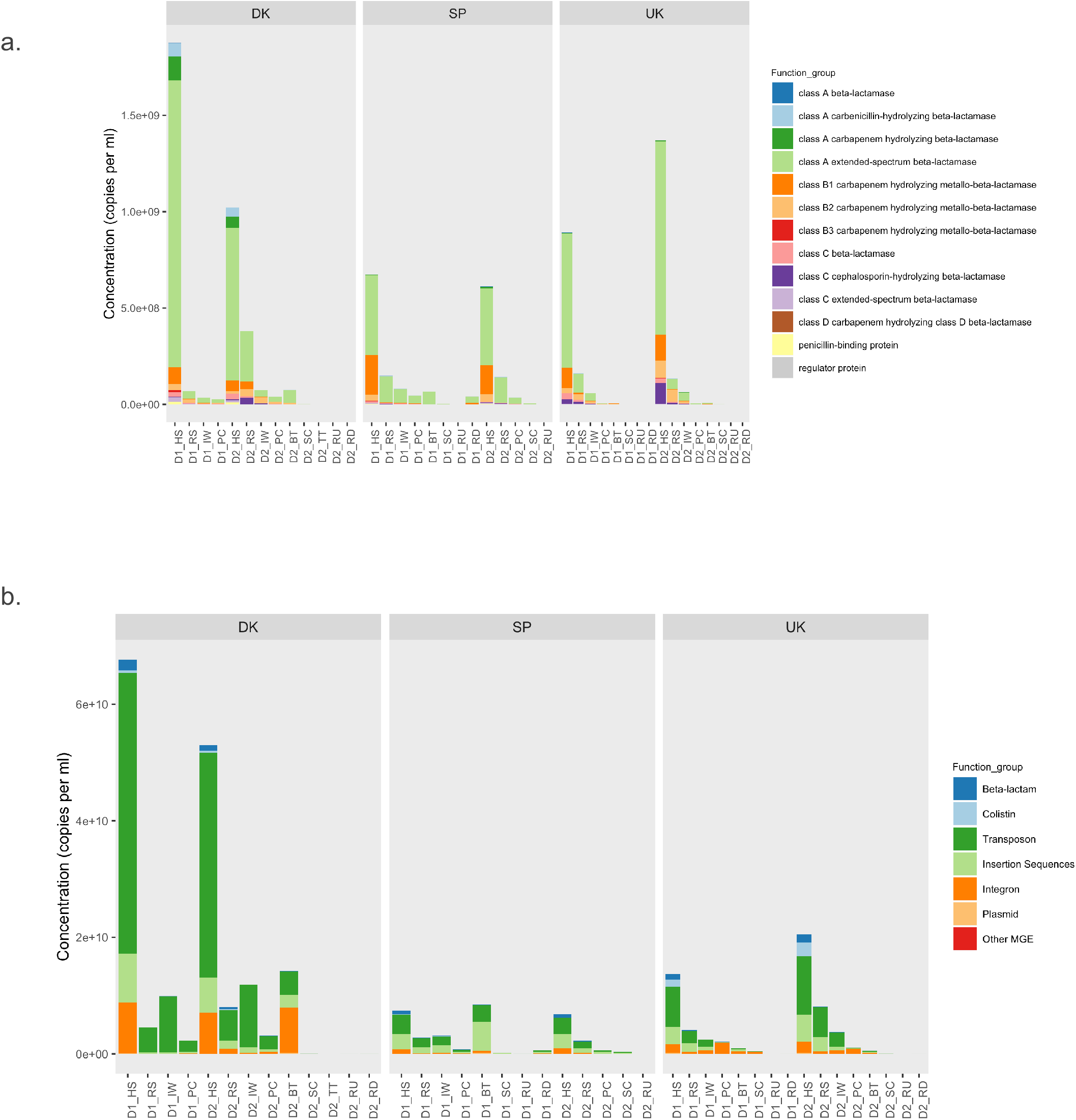
ARG and MGE composition across sampling compartments, days and countries. **a)**. composition profile of ARGs are grouped by function, e.g., β-lactamase genes are classified by their Ambler classification with specific function indicated. **b)**. composition profile of both ARGs and MGEs. ARGs are grouped by the antibiotics they resistant against and MGEs are grouped into insertion sequences, integrons, plasmids, transposons and other MGEs.

A large diversity of ARGs and MGEs was detected across countries; i.e., 79 – 89 out of 118 genes on the qPCR array, including 39 – 47 β-lactam resistance genes and 38 – 41 MGEs. The total concentration of ARGs ranged from 1.6 × 10^7^ to 2.6 × 10^9^ copies per ml, with the highest value found in hospital sewage and the lowest in upstream river samples (Fig. 3a). Among the β-lactamase genes, class A (extended spectrum) and class B2 (carbapenem hydrolysis) were the main groups, accounting for 70% and 16% of concentration across samples on average. Among the class A group, the five dominant genes *cfxA*, *blaBEL*, *blaGES*, *blaVEB* and *blaTEM* comprised on average > 90% of concentration across samples. The main MGE groups were transposon-associated genes like *tnpA-03* and *tnpA-05* (Fig. 3b).

Unlike for the microbial community, ARG and MGE composition was shaped by both country and compartment together (PERMANOVA, both P-values < 0.01) (Fig. S2). For example, although all hospital sewage samples were enriched with diverse ARGs compared with other compartments, distinct dominant genes characterized each country: *mcr-1*, the most abundant ARG (>10%) in hospital sewage in the UK, accounted for less than 1% in DK and SP; *blaVIM*, reaching 2% in SP, was less than 0.1% in DK. Similar country-specific profiles also were apparent for MGEs; e.g., *tnpA-03*, reaching 40% in DK, while only 18% in SP. Genes at low concentration had country and compartment specific occurrence, for example, *blaOXA10,* detected in hospital sewage, residential sewage, influent and primary clarifier in UK, was almost absent in DK and SP; *blaZ*, detected in hospital or residential sewage in SP and UK, was undetected in DK.

Across all UWSs, ARG abundance and diversity gradually decreased along the STP treatment train, and most ARGs were undetectable in the final stage of treatment (e.g., 76 – 84% of β-lactamase genes detected in influent were below detection limit in effluent of secondary clarifier and tertiary treatment) and even those detected were at low concentration (e.g., in DK, 2.0 – 4.8 × 10^3^ copies per ml in tertiary treatment versus 2.8 × 10^6^ – 2.9 × 10^7^ copies per ml in influent). Compared with ARGs, the concentrations of MGEs were more variable despite the consistent decrease from influent to effluent in STP. In contrast to observations in DK and UK, the downstream river in SP was enriched with various ARGs and MGEs, which might be because the plant is under-sized and occasionally discharges sewage without treatment during wet-weather events.

### Hospital sewage has unique ARG profile and high ARG load

In all three countries, ARG and MGE profiles of the hospital and residential sewage differed consistently, even though they had similar microbial community structures (Fig. S1, Fig. 3). The concentration of ARGs in hospital sewage was always more than 5-fold higher than in the residential sewage. In spite of greater abundance disparity, ARG richness was similar, with an average of 38 and 35 ARGs in hospital and residential sewage respectively, and 31 ARGs were shared by the two sewage types. However, the ARG composition clearly differed between the two types of sewage sources; e.g., in the UK, the relative abundance of *mcr-1* was ca. 60% in hospital sewage while less than 10% in residential sewage; in DK, the ARGs of class A β-lactamase of carbenicillin and carbapenem hydrolysis (e.g., *blaCARB*, *blaKPC* and *blaPSE*) was 3 – 5% in hospital sewage while less than 0.1% or even absent from residential sewage. In contrast, the group of class B2 β-lactamase (e.g., *blaCphA* and *blaSFH*) was 0.1% and 1.5% in hospital sewage while 8 – 29% and 18 – 42% in residential sewage in DK and UK respectively. Similar to the ARG profile, the MGE abundance was 3-to 10-fold higher in hospital compared with residential sewage. The estimated hospital contribution to the ARG and MGEs of the total sewage was 10 to 20%, although it contributed less than 3% of the total sewage flow.

We identified 11, 6 and 8 hospital-sewage unique ARGs and MGEs in DK, SP and UK respectively (Fig. 4a). The unique ARGs included β-lactamase genes of class B1 (*cifA*, *blaVIM*, *blaNDM*, *blaSIM*) and class C (*blaFOX*, *blaACC*, *blaACT*, *blaOCH*, *ampC*). In addition, we detected four unique MGEs in hospital sewage (IS5/IS1182 and three plasmid associated elements of IncI, F, HI). We tracked these hospital-sewage unique ARGs and MGEs along the UWSs and several of them were still detected in early STP compartments (i.e., influent and primary clarifier). However, they were undetected in downstream treatment processes, except for IncI in DK, and *blaVIM* in SP and DK. STPs therefore appear effective barriers for the hospital-sewage unique ARGs and MGEs. However, in SP we observed most of the hospital-sewage unique ARG and MGEs in downstream river samples.

**Fig. 4.**
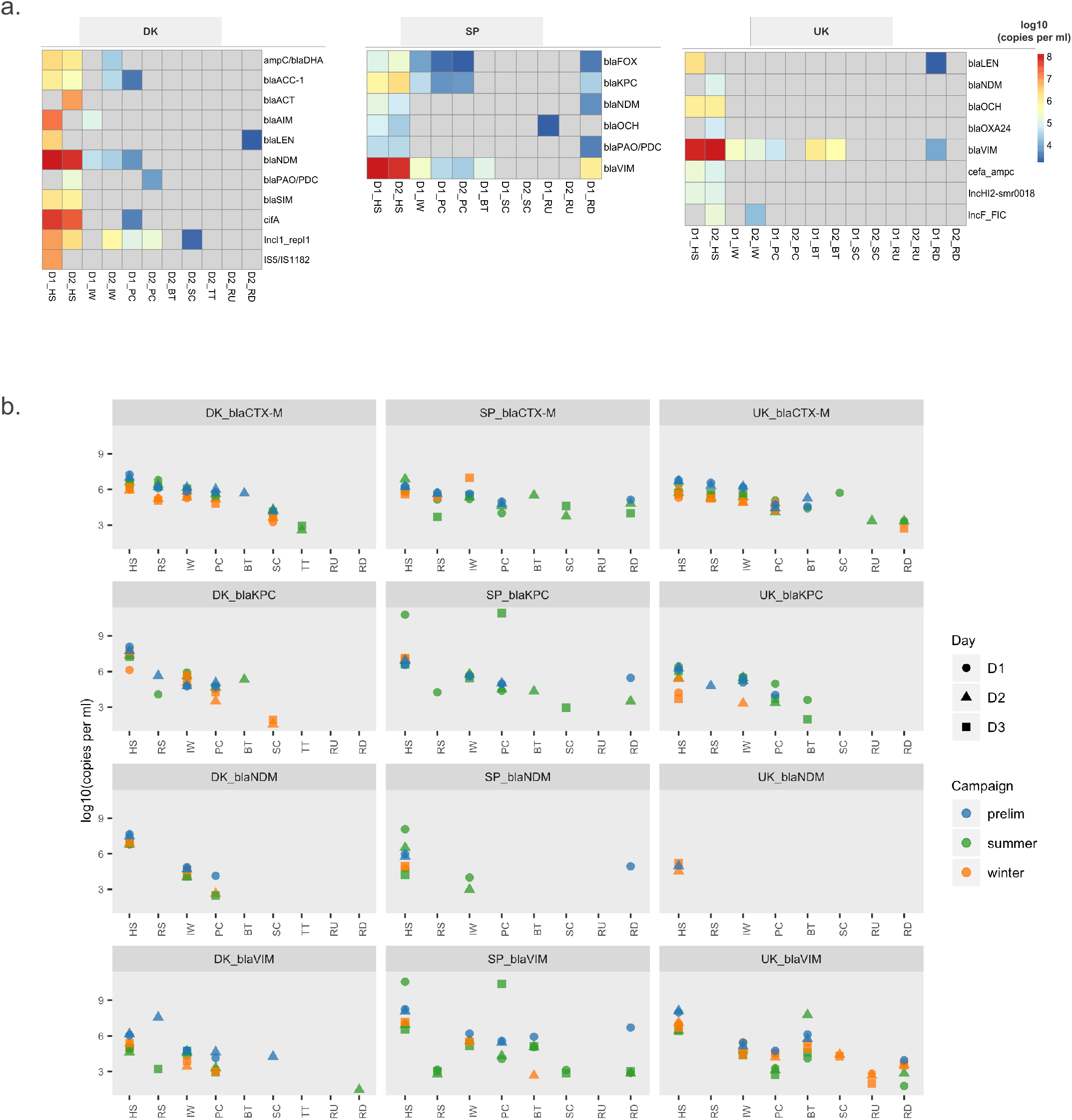
Concentration of hospital unique ARGs along UWS compartment. **a)**. Concentration trend of country-specific hospital unique ARGs along UWS compartments (gray indicates below detection limit). **b)**. concentration profile of four selected ARGs (i.e., *blaCTX-M*, *blaKPC*, *blaNDM* and *blaVIM*) across sampling countries, days and campaigns. Concentration is log10 transformed.

Considering the occurrence of hospital-sewage unique ARGs through the UWS and their clinical importance, four β-lactamase genes – *blaCTX-M*, *blaVIM*, *blaNDM* and *blaKPC* were selected for further validation in the winter and summer campaigns. Besides the three hospital-sewage unique ARGs (*blaVIM*, *blaNDM* and *blaKPC*), we included *blaCTX-M* as an ARG of relatively high abundance across UWS compartments. We observed abundance variation of the four genes among the three campaigns but patterns within each UWS were consistent across seasons (Fig. 4b). STPs were poor barriers for *blaCTX-M* especially in summer. *blaNDM*, the only hospital-sewage unique ARG shared by the three countries, was present in hospital sewage while absent in residential sewage and was readily removed by primary treatment. *blaKPC* and *blaVIM* had the biggest variation among campaigns, and they were even detected in the secondary clarifier or tertiary treatment, as well as in river samples.

### Dynamics of ARG during sewage transport to STP

By coupling the concentration of ARGs, MGEs, and taxa with sewage flow rates, we estimated fluxes entering STP, assuming transport only (i.e., no removal or amplification). This estimated flux was compared with the observed flux in STP influents (Fig. 5, Fig. S3). Among all 322 observed ARGs and MGEs in influent, 243 showed significant removal (with a criterion of 90% or lower than the estimated values). In contrast, 66 ARGs and MGEs appeared to be enriched during sewage transport. Regarding the 74 – 85 ARGs and MGEs that were detected in raw sewage, disappearance in influent was most apparent for 15% – 27% of the ARGs and MGEs, as they were reduced below detection limit.

**Fig. 5.**
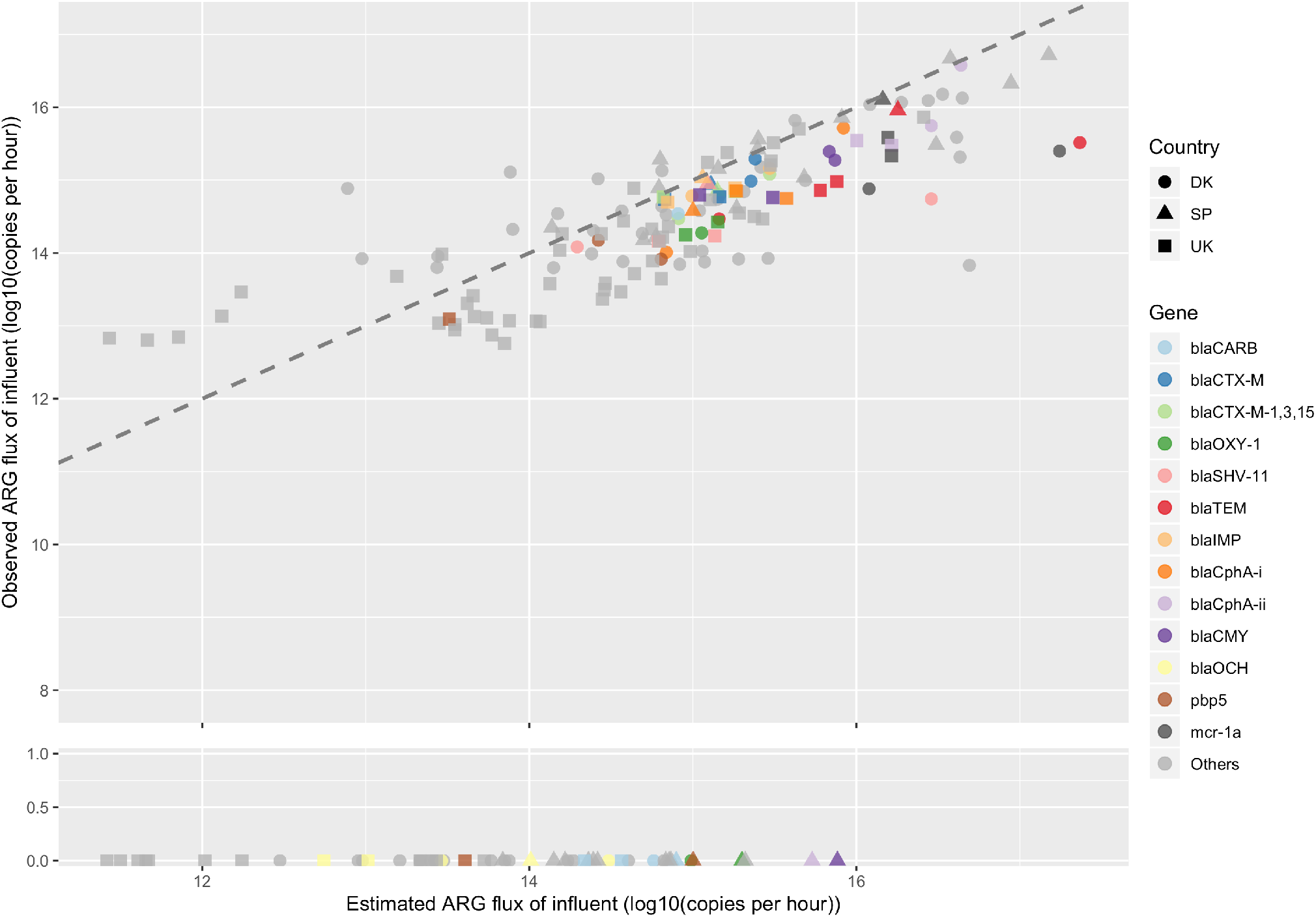
Comparison of estimated and observed ARG flux of STP influent. Points indicate ARGs, in which 13 ARGs decayed consistently across countries are highlighted by color and others are gray. The country in which ARGs are detected is indicated by shape. The gray dash line is reference line of slope = 1, indicating no difference between estimated and observed ARG flux. Flux value is log10 transformed.

Across the three countries, 13 ARGs and 15 MGEs were removed in all cases of sewage transport (Table S2). In particular, *blaOCH* was always below detection limit in the STP influent; *blaOXY-1*, *blaCARB* and *blaTEM* displayed on average > 80% removal. Of the 15 MGEs, there were 9 ISs, 4 transposons, and 2 plasmids of the IncN and IncQ groups. Pairwise comparison between the observed and the estimated flux identified statistically significant removal of 70% across genes, countries and sampling days (Wilcoxon test, P-value < 0.01), indicating that removal was a general phenomenon during sewage transport. In fact, for 36 – 67 ARGs and MGEs, removal in sewer exceeds that occurring within the STP across countries (i.e., average removal ratio 65% by sewer and 34% by STP) (Fig. S4).

One explanation for such substantial ARG removal during sewage transport is dilution to below detection limit of our analytical methods. Indeed, if a gene is only present in hospital sewage, the expected concentration will decrease by more than an order of magnitude as hospital sewage contributes to less than 3% of the total sewage flow. Such dilution to below detection limit however can only explain 14% of the observations. We then tested another possible explanation – whether ARG and MGE removal could be explained by host disappearance. We therefore examined the estimated and observed fluxes of each taxon observed in microbial communities in STP influent (Table S3). Taxa with the highest removal were intestinal types with preference for anaerobic environments like *Clostridriales* (e.g., *Lachnospiraceae*, *Ruminococcaceae*), *Lactobacillales* (e.g., *Enterococcaceae*, *Lactobacillaceae*, *Streptococcaceae*), *Bacteroidales* (e.g., *Prevotellaceae*), and *Enterobacteriales* (e.g., *Enterobacteriaceae*). More than 50% of these intestinal bacteria displayed more than 70% decay, especially members of the *Clostridriales*, *Bacteroidales* and *Enterobacteriales* orders. Since these bacteria typically carry a diversity of ARGs, we cannot exclude the possibility that their removal might cause corresponding ARG loss.

### ARG groups with different persistence in UWSs

It is apparent that not all ARGs and MGEs display the same fate through UWSs, with some restricted to a few compartments, and others more persistent and present in most compartments. We therefore clustered the 118 analyzed ARGs and MGEs into four groups according to their concentration profile across samples (Fig. 6a). These groups with increasing persistence fate through the UWS were named persistence Group I, II, III, IV based on the most downstream location where the genes were still detected: Inlet to STP (Group I), primary clarifier (Group II), biological treatment reactor (Group III), secondary clarifier (Group IV). ARGs and MGEs with occurrence patterns that did not fit any of the four groups were gathered into a category named ‘Other’. The genes in this group were either ARGs with sporadic occurrence, or MGEs prevalent throughout the whole UWS at relatively high abundance.

**Fig. 6.**
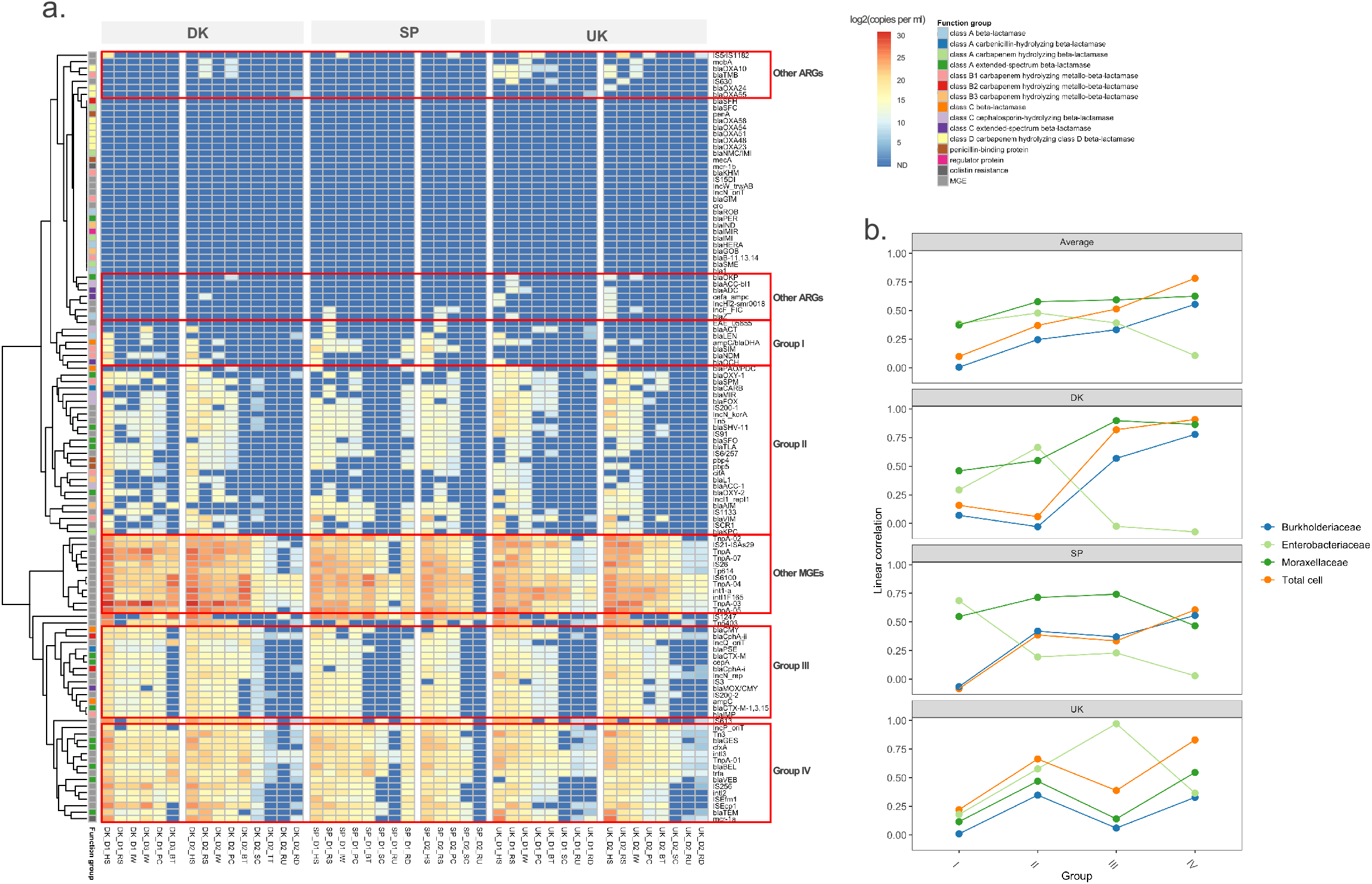
ARG and MGE grouping by concentration profile across all samples, and their correlation with four bacterial groups. **a)**. Heatmap of ARG and MGE concentration along UWS compartments. Samples at x-axis are grouped by country and day along the sequential compartments. ARG and MGE at y-axis are clustered into six groups (Group I, II, III, IV, and other ARGs, other MGEs). Functional annotation of gene is indicated by color bar. Persistence of groups through the UWS does not necessarily correlate with high concentration, as there is no significant impact of concentration level in influent on the grouping (ANOVA, p-value > 0.05). **b)**. Linear correlation of concentration between Group I – IV and four bacterial groups (i.e., *Enterobacteriaceae*, *Moraxellaceae*, *Burkholderiaceae* and total cell). The top panel is the averaged correlation across countries, followed by three panels of each country.

Since the fate of ARGs is mainly determined by their mobility potential and/or by the fate of their hosts, we expect that ARG groups with higher persistence might be associated with higher mobility potential and broader host diversity. We thus examined the relationships between ARG, MGE, and community member persistence through network analysis. We hypothesized non-random co-occurrence pattern among ARGs, MGEs and microbial taxa (i.e., significant positive correlations of Spearman’s R^2^ > 0.8, P < 0.05) to be indicative of ARG mobility potential and ARG-host association. The correlation network consisted of 126 nodes (27 ARGs, 31 MGE, 68 orders) and 1074 edges. The largest group was composed of 20 ARGs and 24 MGEs that strongly associated with 11 taxa (Fig. S5). In particular, ARGs of the two relatively persistent Groups III (e.g., *blaCTX-M*, *blaPSE*, *blaMOX*/*blaCMY*, *ampC*, *cepA*) and IV (e.g., *blaTEM*, *blaGES*, *cfxA*, *mcr-1*) significantly correlated with MGEs (transposon, IS, integron) as well as taxa *Pseudomonadales*, *Enterobacteriales, Clostridiales,* and *β-proteobacteriales*. Compared with Group III and IV, there were only 5 ARGs of Group II significantly correlated with MGEs or taxa, in which *pbp5* and *blaSHV-11* were the only ARGs strongly connected with both MGEs (e.g., transposon and integron) and taxa (e.g., *Enterobacteriales* and *Clostridiales*). In contrast to Group II, III and IV, we did not detect significant correlation between ARGs of Group I and any of MGEs or taxa.

To further identify specific taxa with strong association to ARG persistence, we applied a linear regression model between abundance of the ARG groups and four bacterial groups (Fig. 6b), including the total cell and three taxa at the family level: *Enterobacteriaceae*, *Moraxellaceae*, *Burkholderiaceae.* ARGs of the Group I and II correlated relatively strongly with members of *Enterobacteriaceae* and *Moraxellaceae* (e.g., R^2^ > 0.6 between Group I/II and *Enterobacteriaceae* observed in all the three countries). Group I and II however correlated weakly with total cell numbers. The more persistent Groups III and IV correlated more strongly with *Moraxellaceae* and total cells than the other groups (e.g., R^2^ > 0.6 between Group IV and total cells observed in all the three countries).

## Discussion

Results across the three countries show that the UWS compartment strongly shapes the resident community profile. Such clear effects of compartment on communities were identified in previous studies across UWS^9,10,14,19^. This is likely due to the distinct environmental conditions in each UWS compartment. In the transition from the human gut to the sewer, the enteric microbial community is subject to a strong shift from higher temperature (37ºC) and anaerobic conditions to lower temperature (20 – 25ºC) and moderately anaerobic or aerobic conditions. Such change in conditions not only affects survival of the introduced community but also supports a sewer-specific microbial community – including *Arcobacter, Acinetobacter* and *Aeromonas*^15,16^. These bacteria would significantly influence the outcome of the introduced community through processes like competition and dilution. As a result, we observe a substantial shift of microbial community as sewage is transported from upstream of the conveyance system to the STP. This is evident in the large decrease of enteric bacteria (*Clostridriales*, *Bacteroidales* and *Enterobacterales*) from 37% to 21% on average. This mirrors previous results indicating that only around 20% of the microbes in sewage come from human enteric bacteria^20^.

Upon entering the STP, sewage bacteria experience another series of evident environmental changes, including quiescent sedimentation tanks and highly oxic conditions in the biological reactor. These compartments certainly enrich microorganisms specifically adapted to these unique conditions. In the common biological treatment process of activated sludge, stable resident bacteria have been revealed in global-scale and long-term surveillances, such as bacteria from classes of *Alphaproteobacteria*, *Actinobacteria*, *Gammaproteobacteria*, *Acidimicrobia*, *Sphingobacteria* and *Anaerolineae*^21,22^. Indeed, these bacteria were also the dominant (accounting for 78%) in the biological treatment compartment in STPs of DK and SP, while much less in biofilter system in UK STP and other compartments. The dynamics of UWS microbial community therefore is largely shaped by environmental conditions and resident communities of the compartments they go through.

In parallel with these community shifts, the ARG load in sewage was significantly reduced through the UWS. Here, we quantify the extent of attenuation (65%) across a suite of ARGs during sewage transport. Earlier cultivation-based studies suggested large decreases of resistant bacteria in sewage from immediate discharge to STP entrance^23,24^, and direct monitoring of β-lactamase genes *blaTEM* and *blaKPC* along the length of a sewer pipe also revealed reduction^25^. Such attenuation is presumably caused by the loss of specific ARG host bacteria; the substantial decrease of enteric bacteria during sewage transport reduces the ARGs harbored by those bacterial groups. In addition, since carrying ARG often results in fitness burden^26^, host bacteria of the same ecotype are likely to be outcompeted. This would be especially likely for ARGs carried by plasmids with high fitness cost^27^ and when selective pressures for plasmid carriage are absent^28^.

In contrast, we also observed enrichment of some ARGs during sewage transport, potentially caused by either sources not sampled in this study or gene proliferation by hosts growing in the sewers. The sewer itself houses rich microbial communities in both planktonic and biofilm phase, which have been revealed as potential reservoirs of ARGs like *blaTEM*^25^. Despite selective enrichment, most ARGs not removed during sewage transport can be substantially removed by treatment processes in the STP; on average 26 out of 30 ARGs detected in influent were absent in effluent. The ARG shift driven by sewage transport appears common in UWSs since we observed consistent patterns across countries and sampling campaigns, in spite of the imprecisions associated with measurements of solid concentrations and flow rates in sewers. Given such obvious ARG shifts, ARG profiles in the sewage versus the human gut differ greatly^14,29^. Recently, a plan for global surveillance of antibiotic resistance in sewage was outlined as an affordable way to survey community ARG prevalence^30^. Based on the notable ARG decay during sewer conveyance observed in this study, we argue that simple sewage surveillance will substantially underestimate the real ARG load in the studied population and that rigorous studies are warranted to examine to what extent the ARG profiles in sewage mirror community ARG profiles.

We consistently identified hospital sewage as significant point sources of ARGs in the UWSs across countries and seasons, which has been seen before^10^. In fact, hospital and residential sewage can be very different, which we saw here. For example, ARG concentrations of 2.0 – 3.0 × 10^10^ copies per ml (similar level as in this study) with β-lactamase genes as dominant were observed in hospital sewage in Spain, around 10-fold higher than in residential sewage^10^.

Such relatively high ARG load in hospital sewage was also verified in other EU countries. For example, the Netherlands and France had a 16- and 26-fold higher concentration of β-lactamase genes in hospital versus residential sewage respectively^19,31^, which is even higher than the 10-fold difference observed in our study. In a survey of five hospitals in Singapore^9^, a few variants of β-lactamase genes were only detected in hospital sewage (e.g., *blaIMP*, *blaNDM* and *blaVIM*). Except *blaIMP*, we also identified most of these genes as unique to the hospital in at least one of the three countries (i.e., DK, SP and UK) with *blaVIM* of relatively high abundance in both SP and UK. Consistent with our study, researchers have revealed the complete removal of the hospital-unique β-lactamase genes in STPs, even though a few other persistent genes like *blaVEB* and *blaTEM* were still present in effluent despite remarkable abundance reduction by STP^9^.

We identified ARG groups with distinct persistence across the UWS, which might be associated with their enteric versus environmental origin, or plasmid versus chromosomal location. For example, *blaOXA*, a carbapenemase encoding gene, is commonly found in *Acinetobacter baumannii* clinical isolates colonizing hospitalized patients and is usually located on the chromosome^32,33^. These ARGs are therefore more likely stably maintained within the specific bacterial groups rather than transferred to other environmental bacteria. On the contrary, ARGs of environmental origin, especially those harbored by sewer or STP communities, may well be maintained in a broad range of sewage resident hosts^25,34^. Moreover, compared with many chromosomal-borne β-lactamase genes^35^, MGEs especially the broad-host-range plasmids^36,37^, encoding a wide variety of β-lactamases (e.g., *blaGES*, *blaTEM*, *blaSHV*, *blaIMP*) have become the most prevalent mechanism leading to their global dissemination^38^. Although plasmid carriage often imposes fitness burden on host bacteria, compensatory adaptation can rapidly reduce the cost and improve plasmid persistence^39^. A recent laboratory study has revealed that complex microbial communities have hitherto unrecognized potential in maintaining plasmids by gaining phylotype-level fitness benefits even under non-selective conditions^40^. Additionally, conjugal plasmids carrying ARGs can be stable in multispecies communities through source-sink microbial interactions^41^. Indeed, *blaTEM*, encoding the most common plasmid-mediated β-lactamase^42^ was identified within the high persistence Group IV, and displays significant co-occurrence with various MGEs, whereas *blaLEN* – a typically chromosomal β-lactamase gene^43^, belongs to the low persistence Group I. Given the observed ARG persistence pattern, highly resolved strategies are necessary for efficient ARG containment. Popular approaches for ARG containment have focused on control of enteric bacteria^44^. We argue that elimination of enteric bacteria can only contain part of ARGs (i.e., those of persistence Groups I and II), while community-level strategies are necessary for controlling ARGs of the more persistent group (i.e., Groups III and IV).

Here, we documented that sewage, especially discharged from hospitals, carries substantial loads of ARGs that progressively decline in UWS during conveyance in the sewers. Such attenuation needs to be considered during sewage based antibiotic resistance surveillance. We identified four ARG groups of different persistence fate in the UWS compartments of the three countries. The less persistent groups were associated with putative host taxa like *Enterobacteriaceae* and *Moraxellaceae*, while the more persistent groups appear horizontally transferred. Overall, the attenuation versus persistence of ARGs in UWS revealed in this study can inform management decisions on containment of antibiotic resistance in urban sewage and assist policymakers on the selection of appropriate sampling points for epidemiological ARG surveillance.

## Methods

### Site description, sampling and metadata collection

Three independent sampling campaigns were performed. The first lasted two days in spring/summer 2017; the second, three days in winter 2018; and the last, three days in summer 2018. At least 1.5 l 24-hr flow proportional, refrigerated samples were collected from each sampling point, except in SP where 10-hr composite samples (from 8:00 to 18:00, taken every 2 hours) were collected, and for the activated sludge, a grab sample was obtained at around 11:30. Due to their low biomass content, at least 20 l of STP effluent and the river water were collected and concentrated, using dead-end ultra-filtration with a dialysis filter Rexeed 25A (Asahi Kasei, Chiyoda, Japan) as described elsewhere^45^. For molecular analysis, 100 ml of 10-time concentrated biomass suspension was stored frozen at −80°C in 20% glycerol prior to DNA extraction

### DNA extraction

A volume of 500 μl of thawed sample was used for DNA extraction with NucleoSpin Soil kit (MACHEREY-NAGEL, Düren, Germany) following the user manual using SL2 lysis buffer. Bead-beating step was performed on a FastPrep-24 Classic homogenizer (MP Biomedicals, Irvine, CA, USA) for 30 seconds at 5 m/s. The rest of the extraction procedure was performed on an EpMotion 5075 liquid handling platform (Eppendorf, Hamburg, Germany). Finally, DNA was eluted in 100 μL of preheated (68°C) Buffer SE and stored at −20°C. All DNA extracts were quantified using a Qubit^®^ Fluorometer and HS dsDNA kit (Invitrogen, Maryland, MD, USA).

### High-throughput qPCR array

Primers used in the 120-assay setup were a subset of the ARG 2.0 panel (103 primers) with the addition of 17 new qPCR primer sets. New primers added to the high-throughput qPCR array subset as part of this work included: *blaAIM*, *blaGIM*, *blaKHM*, *mcr-1*, *blaNMC/IMI*, *blaOXA23*, *blaOXA24*, *blaOXA48*, *blaOXA51*, *blaOXA54*, *blaOXA55*, *blaOXA58*, *blaSFC*, *blaSFH*, *blaSIM*, *blaSPM-45*, and *blaTMB*. Reference sequences for these genes were obtained from ARG-ANNOT^46^, MEGARes^47^, and Genbank. Sequences of high similarity were obtained using BLAST. Following collection and curation, sequences were aligned using MEGA and primers were designed using RDP EcoFunPrimer design tool (https://github.com/rdpstaff/EcoFunPrimer)^48^. Specificity of primer candidates was evaluated using BLAST and selected primers were chosen based on targeted gene coverage, specificity, and thermodynamic properties. The high-throughput qPCR analysis was conducted on Takara’s SmartChip qPCR platform which allows analysis up to 5,184 100 nl reactions simultaneously (120 assays with 14 samples; with three technical replicates). Samples and primers were dispensed in loading plates per manufacturer protocols with LightCycler® 480 SYBR Green I Master Mix and placed in the chip-loading system which loads the high-density chips automatically, with three technical replicates per sample-primer combination. A cycle quantification value of 28 was used as a cut-off threshold for analysis of positive events, and only assays with amplification in a minimum of two technical replicates were included in downstream analysis. Copy numbers were estimated using a previously described equation^49^, and relative abundance of a target gene was determined as the ratio of average target gene to 16S rRNA gene copies. In further data analysis, β-lactamase gene were grouped by Ambler classification with functions specified.

### 16SrRNA sequencing and analysis

Bacteria and Archaea community compositions were assessed by high-throughput amplicon sequencing. A fragment spanning the hypervariable regions V3-V4 of the 16S rRNA gene was amplified from each DNA sample by an initial PCR step using primers Uni341F (5’-CCTAYGGGRBGCASCAG-3’) and Uni806R (5’-GGACTACNNGGGTATCTAAT-3’) originally published by Yu et al.^50^ and modified as described by Sundberg et al.^51^. In a second PCR reaction step, the primers additionally included Illumina specific sequencing adapters and a unique dual index combination for each sample. After each PCR reaction, amplicon products were purified using HighPrep™ PCR Clean Up System (AC-60500, MagBio Genomics Inc., USA) paramagnetic beads using a 0.65:1 (beads:PCR reaction) volumetric ratio to remove DNA fragments below 100 bp in size and primers. Samples were normalized using SequalPrep Normalization Plate (96) Kit (Invitrogen) and pooled using a 5 μl volume for each sample. The pooled samples library was concentrated using DNA Clean and Concentrator™-5 kit (Zymo Research, Irvine, CA, USA). The pooled library concentration was determined using the Quant-iT™ High-Sensitivity DNA Assay Kit (Life Technologies). Before library denaturation and sequencing, the final pool concentration was adjusted to 4 nM before library denaturation and loading. Amplicon libraries sequencing was performed on an Illumina MiSeq platform using Reagents Kit v3 [2 × 300 cycles] in paired-end mode. Raw sequence reads were trimmed of primer sequences used in first PCR, discarding read pairs for which any of the two primers sequences could not be detected using cutadapt version 2.3^52^. Primers-trimmed sequence reads were error-corrected, merged and amplicon sequence variants (ASVs) identified using DADA2 version 1.10.0^53^ plugin for QIIME2^54^ with the following parameters: truncL = 230, truncR = 215; trimL=8, trimR=8 and otherwise defaults parameters. A separate denoising was performed for each run as recommended by DADA2. Each ASVs was given a taxonomic annotation using *q2-feature-classifier* classify-sklearn module trained with SILVA SSU rel. 132 NR99 database^55^. Prior to training the classifier, the V3-V4 region of SILVA database sequences was extracted at the same primers position. Raw sequence data are publicly available at the NCBI-SRA database under BioProject accession number PRJNA672724.

### qPCR detection and analysis

*blaVIM, blaNDM* and *blaKPC* were quantified using the SsoAdvanced™ Universal Probes Supermix (Bio-Rad), employing the following thermocycle program: (i) 3 min of initial denaturation at 95 ºC, and 40 cycles of (ii) 5 s denaturation at 95 ºC, and (iii) 30 s annealing/extension at 60 ºC. In addition, qPCR also was used to quantify *blaCTX-M* and total eubacteria using a SYBR green-based method assay. SYBR-green reactions were conducted using SsoAdvanced™ Universal SYBR® Green Supermix (BioRad), employing the following thermocycle program: (i) 2 min of initial denaturation at 98ºC, and 40 cycles of (ii) 5 s denaturation at 98ºC, and (iii) 5 s annealing/extension at 60 ºC. Primer sets for qPCR assay were listed in Table S4. All assays were done in triplicate using the BioRad CFX C1000System (BioRad, Hercules, CA USA). In order to avoid inhibitor effects, DNA samples were diluted to a working solution of 2 ng/ul and an internal control DNA was used in SYBR-green reactions. Standard curves for each set of primers were constructed using plasmid clones of the target sequences of between 1.0 × 10^2^ and 1.0 × 10^8^ copy numbers, which were used in triplicate and in parallel with each qPCR run.

### Concentration and flux calculation for taxa and genes

The cell concentration N (cells l^−1^) in each sample was derived from the VSS measurement (mg l^−1^). It was calculated using Eq. (1)^56^, assuming a bacterial biovolume (V) of 0.25 μm^3^ cell^−1^, a carbon content per unit of cell volume (Cs) of 310 fg C μm^−3^ and considering that only ~53% of a cell’s dry weight constitutes carbon.

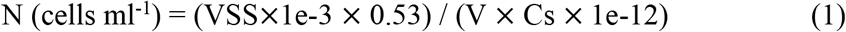

The cell concentration of each taxon Nt (cell ml^−1^) in the sample was then obtained by multiplying the total cell concentration with the relative abundance of the taxon in the amplicon library.

The concentration of each ARG or MGE (copies ml^−1^) was calculated using Eq. (2) by considering the cell concentration N (cells ml^−1^), the average 16S rRNA gene copies per cell (16Scell, copies cell^−1^) and relative abundance of ARG or MGE (RA, copies (16S rRNA gene copy)^−1^). For each amplicon library, universal 16S rRNA gene amplicon sequencing data was used to perform CaRcone analysis to obtain the average 16S rRNA gene copies per cell (R script https://github.com/ardagulay/CaRcone-Community-average-rRNA-gene-copy-nrestimator)^57^.

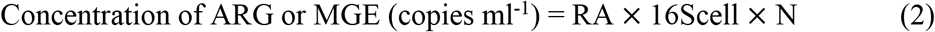

The observed flux of each gene or taxon (copies hour^−1^ or cells hour^−1^) at the STP entrance was obtained by multiplying the concentration of ARG or MGE (copies ml^−1^ or cell ml^−1^) with the average flow rate (ml hour^−1^) of the sewage entering STP. This was compared with the expected flux calculated from the abundances measured in the hospital and residential sewers, under the assumption of simple conveyance of the genes or the taxa from the sewer sampling points to the entrance of STP, and taking into account the respective contribution of both sources to the STP influent. Deviation from the expected flux indicates violation of the ‘conveyance only’ assumption, for example because some taxa are outcompeted by other, resulting in their effective removal. Pairwise comparison between the two fluxes was performed by Wilcoxon test, in which only the ARGs or MGEs above detection limit in qPCR array were included. For each gene, the relative contributions of sewer conveyance and treatment in STP to total removal were calculated respectively as (F1-F2)/F1 and (F2-F3)/F1, where F1 is the flux entering the sewers, including hospital and residential sewage; F2, the flux entering the STP (i.e., STP influent); and F3 the flux leaving the STP (i.e., effluent).

### Statistical analysis

Non-metric multidimensional scaling (NMDS) was carried out using Bray-Curtis dissimilarity matrices that were based on concentration of taxa or genes. Permutational multivariate analysis of variance (PERMANOVA) tests were conducted to evaluate the effect of the country, compartment and campaign on microbial community and gene profile, since it is robust to deviation from parametric assumptions. ARG&MGE grouping was performed by hierarchical cluster analysis based on distance measure of Pearson correlation. Network analysis was based on the correlation matrix developed by calculating all possible pairwise Spearman’s rank correlations among bacterial taxa, ARGs, and MGEs. A correlation between two items was considered statistically robust if the Spearman’s correlation coefficient (R^2^) was ≥ 0.8 and the P-value was ≤ 0.05. To reduce the chances of obtaining false positive results, P-values were adjusted with a multiple testing correction using the Benjamini-Hochberg method. The robust pairwise correlations formed co-occurrence networks. Network analyses were performed in R, and was visualized and explored to identify its topological properties (i.e., clustering coefficient, shortest average path length, and modularity) in Gephi^58^. Linear regression models were applied to estimate the association between abundance of the ARG&MGE groups and four specific bacterial groups (i.e., *Enterobacteriaceae, Moraxellaceae, Burkholderiaceae* and total cells). The bacterial groups were chosen based on the network correlation with ARG&MGEs, and their high abundance in UWS compartments specifically or generally, e.g., *Enterobacteriaceae* mainly occurring in raw sewage, while *Moraxellaceae* and *Burkholderiaceae* through most UWS. In addition, we included the total cell in the linear regression analysis, assuming ARG&MGEs hosted by multiple taxa will correlate more with the total cell abundance than with that of any specific taxa.

## Acknowledgements

This work was supported by a Joint Programming Initiative-Antimicrobial Resistance grant (JPI-AMR; DARWIN project 7044-00004B). We appreciate the outstanding help of the staff at VandCenter Syd (Odense, Denmark), Viaqua (Santiago de Compostela, Spain), and Northumbrian Water Limited (Durham, UK) in accessing and sampling their systems.

## Author contributions

L.L., J.N., M.Q., A.D., and B.F.S. designed the study. M.Q., A.D., and S.B. performed the sampling. Z.Y. extracted DNA. S.Y. and M.R.W. performed HT-qPCR. M.Q. performed qPCR. L.L. and J.N. did all the amplicon sequence analyses. L.L., A.D., and B.F.S. interpreted the results. L.L. drafted the manuscript with input from A.D., B.F.S., S.J.S., J.N., D.W.G., J.L.R. All authors contributed to manuscript revisions and have read and approved the final version of the manuscript.

## Data availability

The sequence data analyzed in this study are available in NCBI-SRA database under BioProject accession number PRJNA672724.

