## Supporting information for "Extended spectrum β-lactamase and carbapenemase genes are substantially and sequentially reduced during conveyance and treatment of urban sewage"

**Table S1a**. Information on the catchments

| City | Approx. population in sampled community sewer | Hospital beds | WWTP population equivalent |
|---|---|---|---|
| Durham (UK) | | ~ 450 | 35,000 |
| Odense (DK) | 14,200 | ~ 1,000 (http://ouh.dk/wm397745) | Capacity: 385,000<br>Actual load (2018): 235,000 |
| Santiago de compostella (SP) | 9,600 | 1,395 (https://www.mscbs.gob.es/ciudadanos/centros.do?metodo=realizarDetalle&tipo=hospital&numero=150200) | Capacity: 103,000<br>Actual load (2018): 174,500 |

**Table S1b**. Approx. GPS coordinate of sampling points

| | HS | RS | STP | RU | RD |
|---|---|---|---|---|---|
| Durham (UK) | 54°47'22.2"N 1°35'31.5"W | 54°47'35.2"N 1°34'38.1"W | 54°46'58.6"N 1°34'10.1"W | 54°47'01.6"N 1°33'50.6"W | 54°47'09.0"N 1°33'54.1"W |
| Odense (DK) | 55°23'04.8"N 10°22'21.6"E | 55°24'24.9"N 10°23'15.3"E | 55°23'57.8"N 10°25'01.8"E | 55.398930, 10.419135 | 55°24'00.2"N 10°25'23.9"E |
| Santiago de compostella (SP) | 42°87'01.0"N 8°56'93.2" W | 42°88'11.2"N 8°52'63.0" W | 42°87'03.2"N 8°59'79.8" W | 42°87'12.8"N 8°59'48.5" W | 42°86'92.8"N 8°60'06.7" W |

**Table S1c**. Dates of sampling

| | 2017 | Winter 2018 | Summer 2018 |
|---|---|---|---|
| Durham (UK) | 23/06 and 06/07 | 13/03; 15/03; and 20/03 | 18/09;25/09; and 28/09 |
| Odense (DK) | 31/05 and 01/06 | 30/01; 20/02; and 21/02 | 12/09; 13/09; and 18/09 |
| Santiago de compostella (SP) | 24/10 | 19/02; 21/02; and 23/02 | 17/10; 19/10; 21/10 |

**Additional Information on sampling**

To avoid major temporal variation in resident populations, and because our catchments have a sizable student population, we sampled outside of the main summer/winter vacations and did not sample on weekend (no sampling between Friday noon to Monday noon). We sampled in dry weather only (less than 2 mm precipitation per day).

**Table S4.** Primer sets for qPCR

| Target | Primer/probe | Sequence (5′–3′) | Amplicon size (bp) | Annealing (°C) | Reference |
|---|---|---|---|---|---|
| *blaCTX-M* | ctx_f | CTATGGCACCACCAACGATA | 103 | 60 | (1,2) |
| | ctx_r | ACGGCTTTCTGCCTTAGGTT | | | |
| 16S rRNA | 1055_f | ATGGCTGTCGTCAGCT | 337 | 60 | (3) |
| | 1392_r | ACGGGCGGTGTGTAC | | | |
| Internal control | gfp_f | TCGGTTATGGTGTTCAATGC | 146 | 60 | (4) |
| | gfp_r | GACTTCAGCACGTGTCTTGTAG | | | |
| *blaNDM* | ndm_f | GCAAATGGAAACTGGCGACC | 275 | 60 | (5) |
| | ndm_r | TACCGCCCATCTTGTCCTGA | | | |
| | ndm_p | FAM-TCGCACCGAATGTCTGGCAGCACA-BHQ1 | | | |
| *blaVIM* | vim_f | GATTGATACAGCGTGGGGTG | 167 | 60 | |
| | vim_r | ACGGYGATGCGTACGTTGC | | | |
| | vim_p | HEX-GACGCGGTCGTCATGAAAGTGCGT-BHQ1 | | | |
| *blaKPC* | kpc_f | CACTGTGCAGCTCATTCAAGG | 270 | 60 | |
| | kpc_r | CGCCGATAGAGCGCATGAA | | | |
| | kpc_p | TxRed-CTGCCGCTGTGCTGGCTCGCA-BHQ | | | |

Figure S1. NMDS profile of microbial community composition of all samples (stress value = 0.10).

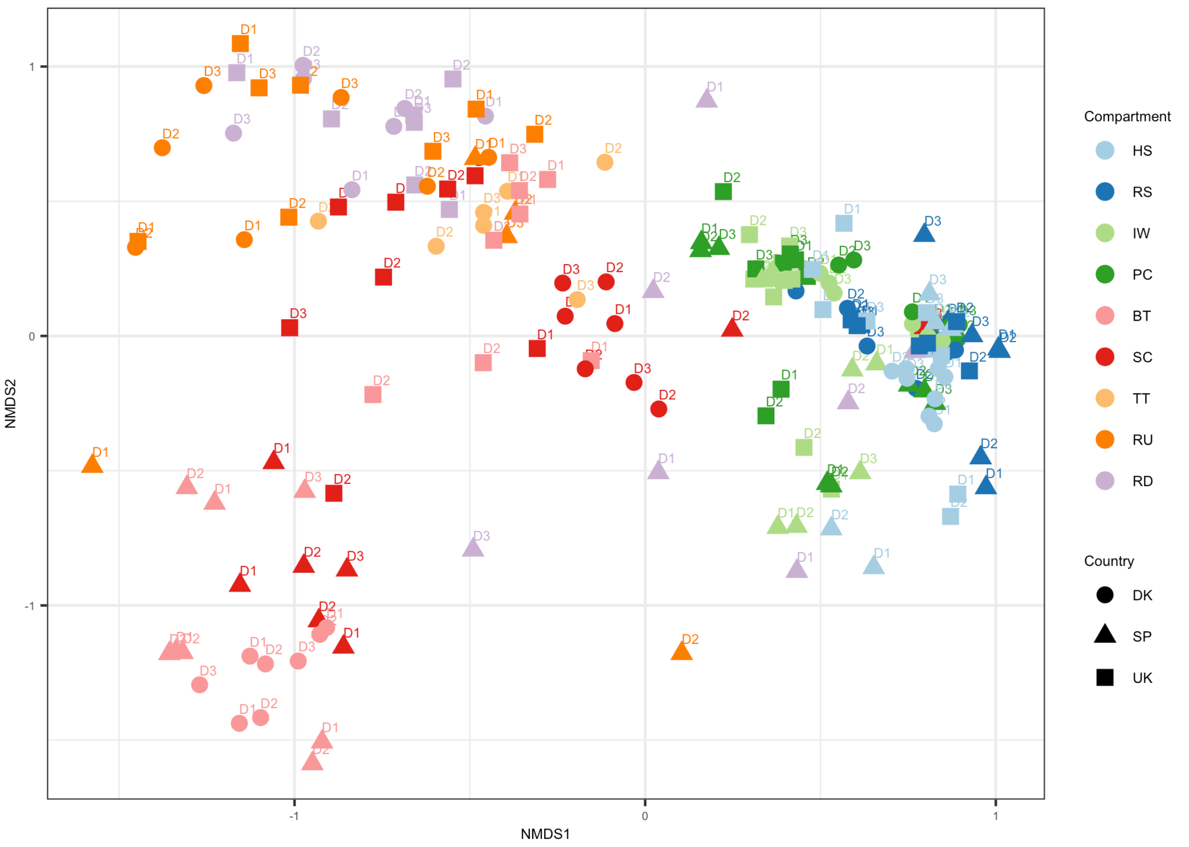

Figure S2. NMDS profile of ARG composition of all samples (stress value = 0.11).

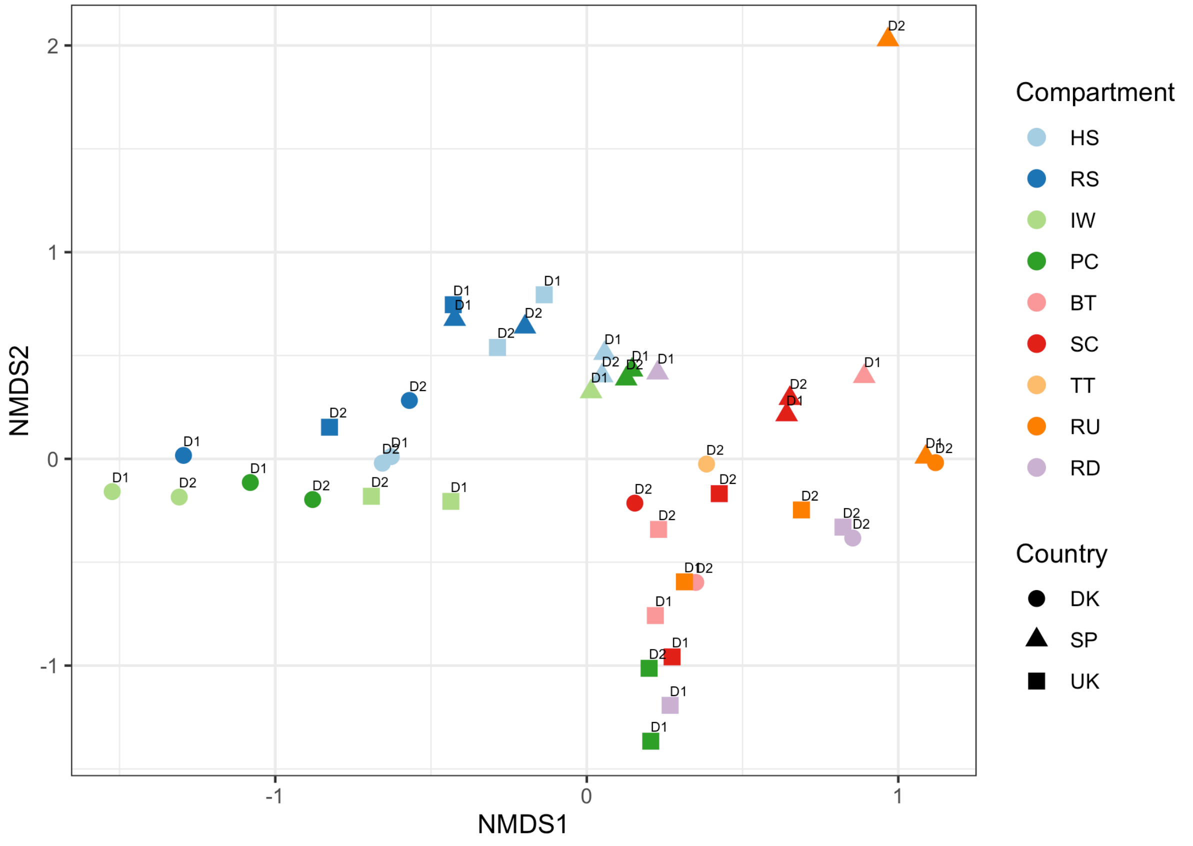

Figure S3. Estimated and observed ARG flux of STP influent. Concentration was log10 transformed.

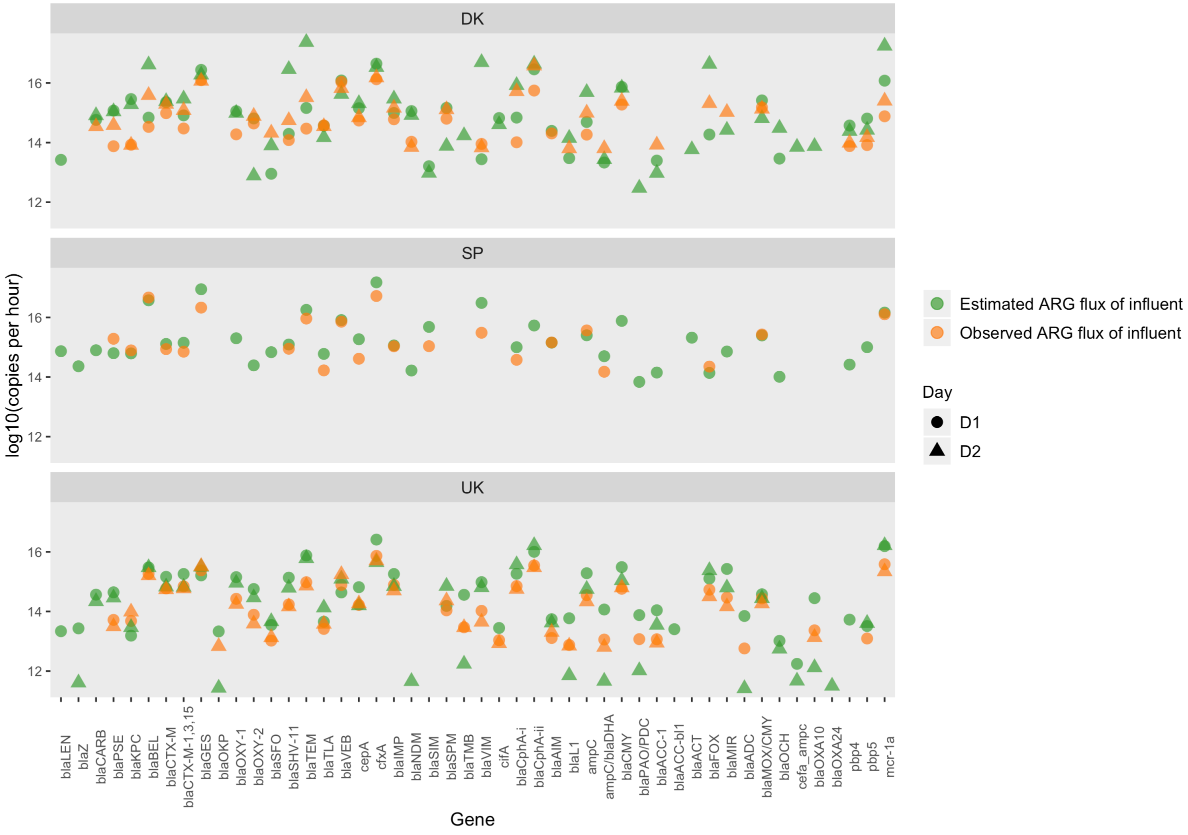

Figure S4. Removal ratio of ARGs by sewage transport and STP processes.

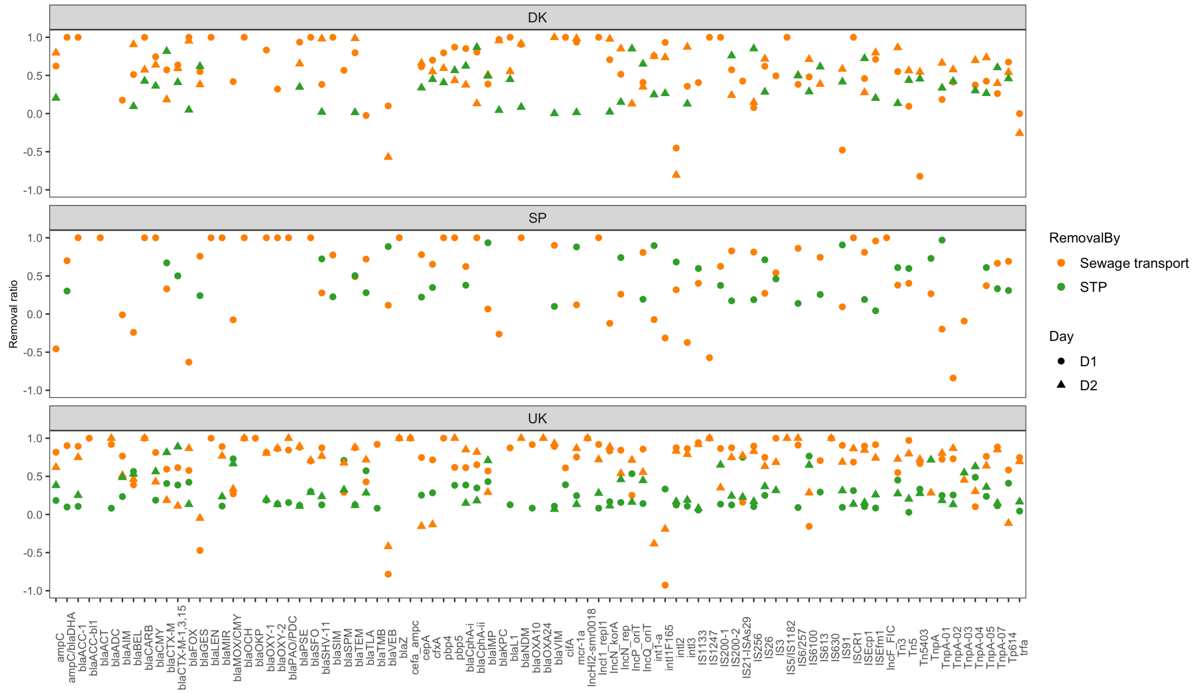

Figure S5. Network analysis revealing co-occurrence patterns among ARGs, taxa (order level), and MGEs. A connection represents a strong correlation (Spearman's correlation coefficient (R2) ≥ 0.8 and P-value ≤ 0.05). ARG groups, taxa, and MGEs are indicated by color.

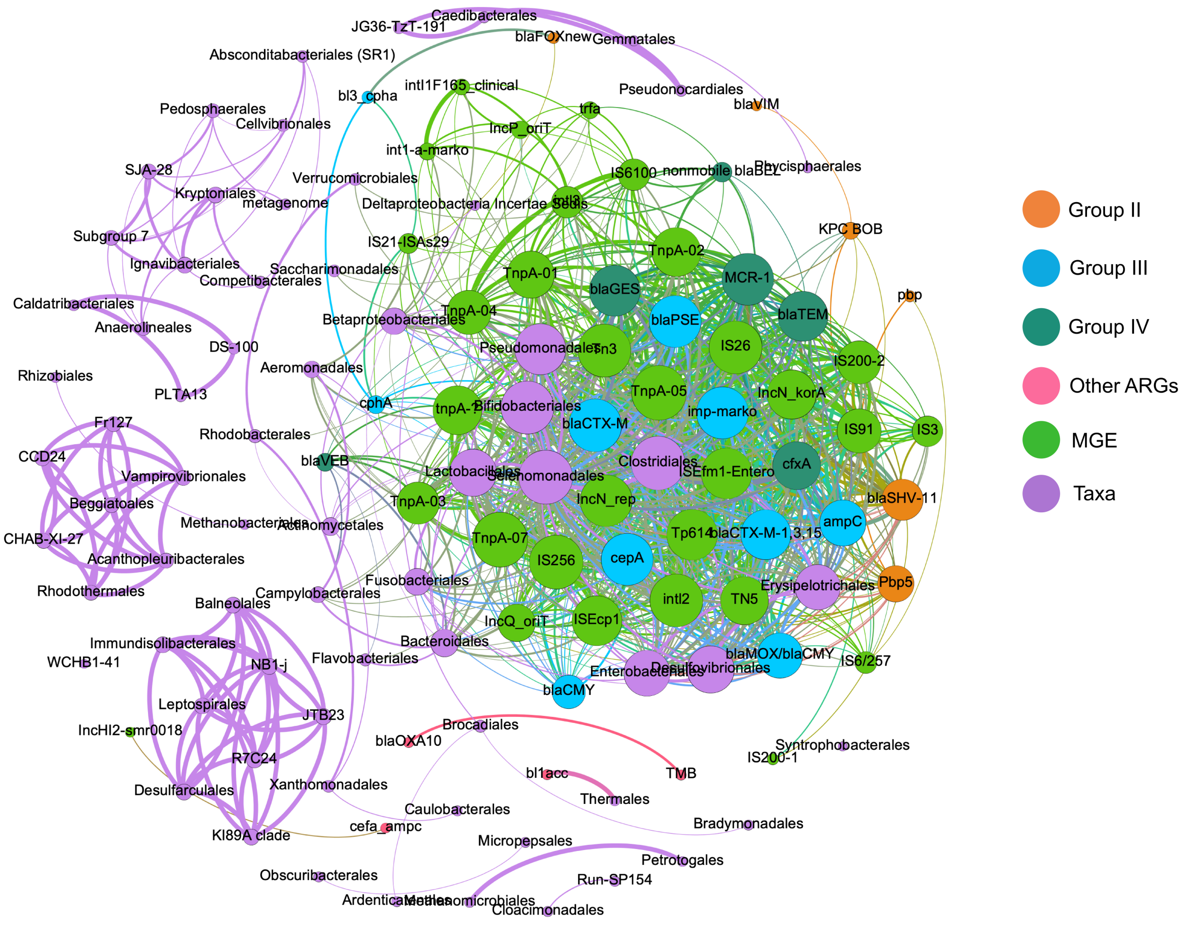
